# Genome-wide investigation of outer membrane protein families under mosaic evolution in *Escherichia coli*

**DOI:** 10.1101/2024.10.30.621157

**Authors:** Xin Cao, Cuihua Cao, Zefan Chen, Jialin Li, Yidong Zheng, Jinjin Wu, Zeling Li, Yueming Hu, Gaofeng Hao, Guoqiang Zhu, Wolfgang Köster, Aaron P. White, Yejun Wang

## Abstract

Several genes in Gram-negative bacteria encoding outer membrane proteins (OMPs) have been reported to show patterns of mosaic evolution featured by mixture of negative selection and local recombination. Here, we proposed a strategy, and applied it to screen OMPs under mosaic evolution in *Escherichia coli* in the genome level. In total, 21 OMP families, including 16 new ones, were detected with the typical patterns of mosaic evolution. An abosolute majority of the protein families are conserved in *E. coli* for the composition, genomic loci and the overall structures. Highly Variable Regions (HVRs) can be recognized, which are frequently located extracellularly within the protruding loops. There are only limited number of major HVR sequence types, within which positively selected sites can be detected ocassionally. Based on the simulated results of multiple models, the OMPs under mosaic evolution are often with good antigenicity, with HVRs of various sequence types coinciding with the B-cell epitopes of the strongest immunogenicity. The study futher broadened our understanding of the characteristics of mosaic evolution and the functions of OMPs in Gram-negative bacteria, laying an important foundation for their potential translational applications.

**Importance:** It is important to understand the evolutionary mechanisms of bacterial OMP-encoding genes, which would facilitate the development of anti-bacterial reagents. This study made the first genome-wide screening of bacterial OMPs under mosaic evolution, and increased the list of candidate OMP families by 3 folds in *E. coli*, far more than we expected. The study further confirmed the hypothesis about the evolutionary, micro-evolutionary and structural features of these OMPs, and facilitated the functional theory of mosaic evolution. Moreover, the findings of limited HVR sequence types and strong immunogenicity of HVRs paved important foundation for application of these OMPs and their HVRs in development of antibodies or other anti-bacterial treatment.

## Introduction

Anti-Microbial Resistance (AMR) poses a severe threat to human health [1]. Globally, drug-resistant bacterial infections claim 1.27 million deaths annually [2]. In China, 145,000 people die due to bacterial AMR each year [3]. It is urgent to develop new strategies to cope with bacterial AMR. On the one hand, there is a need to accelerate the development of novel antibiotics [4–7], therapeutic monoclonal antibodies [8], and other treatment methods [9], such as bacteriophage therapy [10]. On the other hand, it is also crucial to deepen our understanding of the mechanisms of bacterial resistance from the perspectives of bacterial evolution and selection [11, 12], and to improve the sensitivity of bacteria to various existing drugs and treatment protocols [13].

The complicated cell envelope structure of diderm Gram-negative bacteria makes it difficult for extracellular molecules to enter the bacterial cells, posing particular challenges for drug development. To enter diderm bacterial cells, antibiotics first need to cross the outer membrane, and this process is frequently mediated by the bacterial outer membrane proteins (OMPs) [9, 14]. Porins, such as OmpF and OmpC, form hollow channels in the bacterial outer membrane, allowing small hydrophilic molecules to pass through by passive transport, which is the primary route for many antibiotics to enter the bacterial cells [14]. Bacteria can also resist from the antibiotics by reducing their intake through various mechanisms to influence these porins. For example, mutations in the *ompK36* gene of the clinical isolate ST258 of *Klebsiella pneumoniae* result in the insertion of two amino acids into the L3 loop of the protein, significantly narrowing the pore diameter and causing resistance to meropenem [15]. In clinical isolates of *Escherichia coli*, *ompC* mutations have been found to alter the lateral electric field within the pore, inhibiting the entry of cefotaxime, imipenem, and gentamicin [16]. *E. coli* can also reduce antibiotic intake by downregulating *ompC* gene expression [17]. Some porins (e.g., LamB [14]) and other OMPs (e.g., FhuA [18]) bind specifically to antibiotic molecules and transport them passively or actively. Mutations in these genes may also lead to resistance [14, 18]. Besides modification of antibiotic intake pathways, bacteria can also employ other mechanisms to induce resistance, such as upregulating the Resistance-Nodulation-cell Division (RND) efflux pump system to expel antibiotics taken into the cells. Among the three components of the RND system, there is also a β-barrel OMP [19].

In addition to antibiotics, the extracellular portions of bacterial OMPs often serve as target molecules for antibacterial factors such as immune factors and bacteriophages. Therefore, in recent years, some novel bacterial treatment methods, such as therapeutic monoclonal antibodies [20–22] and phage therapies [23, 24], often target bacterial OMPs. Bacteria themselves have also evolved various strategies, such as altering the sequences and structures of the OMPs or downregulating their expression, to evade the attacks of these naturally occurring or artificially designed antibacterial factors, resulting in phenotypes such as immune evasion and bacteriophage tolerance [25, 26].

Exploring the evolutionary characteristics and patterns of bacterial OMPs can aid in designing more effective antibacterial agents and in blocking or delaying bacterial resistance. Evolutionary studies indicate that bacterial OMP genes, such as *fhuA*, *lamB*, *ompA*, *ompC*, *ompF*, and others, are often under positive selection [27, 28], possibly due to their frequent targeting by bacteriophages, immune factors, and other disfavoring factors in the environment [29, 30]. In the laboratory, dynamic sequence variation in *lamB*, *ompA*, and *ompF* can be directly detected by exposing *E. coli* to phage infestation, confirming large evolutionary rates of these OMP genes [11]. These evolutionary characteristics of OMP genes allow bacteria to rapidly derive and select clones tolerant to adverse factors in the current environment, promoting population survival. These evolutionary traits of bacterial OMPs also accelerate the failure of antibiotics, vaccines, and other treatments, thereby increasing the difficulty of antibacterial therapy [11, 31].

Previously, we found that the *fhuA* gene exhibits significant intra- or inter-species local recombination in *E. coli* and *Salmonella* [29]. The local recombination regions, also known as Highly Variable Regions (HVRs), are generally concentrated. FhuA serves as a binding receptor for multiple bacteriophages, and its HVRs are highly consistent with the binding sites of bacteriophages, suggesting that *fhuA* avoids bacteriophage invasion through local recombination [29]. Within the population of *E. coli* with the same HVR type, there is also a certain degree of positive selection within the HVRs [27–29]. Apart from the HVRs, other sequences of FhuA are highly conserved, displaying purifying selection; correspondingly, the overall structure of FhuA proteins across different HVR sequence types is similar, which may be related to the maintenance of the basic function of FhuA, i.e., transporting iron chelates. Subsequently, we further disclosed multiple porin genes in *E. coli*, including *lamB*, *ompA*, *ompC* and *ompF*, which exhibit the typical patterns of mosaic evolution similar to *fhuA*, characterized by the coexistence of local recombination, positive selection, and negative selection [30]. There are OMP genes in other bacteria which were reported to show similar evolutionary patterns [32–35]. Based on these observations, we proposed the mosaic evolution theory of bacterial OMPs. Briefly, bacteria maintain the overall protein structure and function through the conservation of genes and overall sequences, while promoting resistance to bacteriophage and other adverse factors through local recombination and positive selection [29, 30].

To further analyze the distribution of mosaic evolution in bacterial OMPs, in this study, we improved the analytical methods to make genome-wide screening of the OMPs under mosaic evolution in *E. coli*. For the candidate proteins, we further delineated and charicterized the HVRs, observed the HVR sequence types, distribution and micro-evolution, and analyzed their structures and immunogenicity.

## Results

### 1. Mosaic evolution of OMP-encoding genes in *E. coli*

We predicted 183 OMPs from the 4,298 genome-derived proteins of E. coli strain MG1655, and identified their orthologs from other E. coli strains and E. fergusonii ATCC_35469 (Fig. 1A). With RDP4 (version 4.101), we found that 89 OMP-encoding gene families experienced ingenic local recombination (Fig. 1A). Phylogenetic trees were built and manually examined for these genes and the proteins, to further screen those incongruent with the lineage tree for both types of sequences. Exchanging tests were also performed to confirm the local ingenic recombination for each of the gene and protein families. The procedure detected 27 genes showing typical mosaic evolution patterns at both nucleotide and amino acid levels (Fig. 1A).

**Figure 1.**
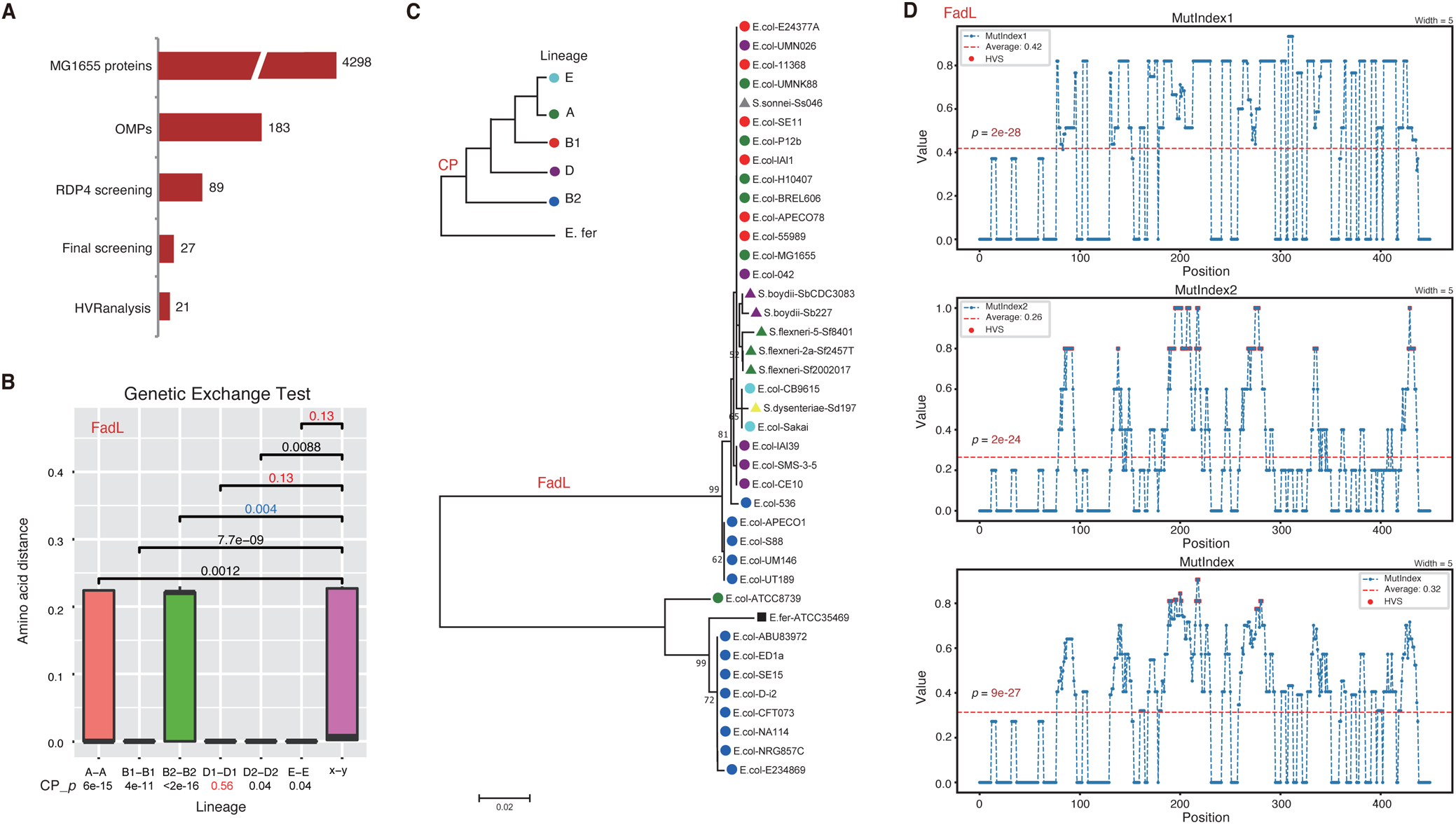
Screening of *E. coli* OMPs under mosaic evolution. **(A)** The OMPs and candidates under mosaic evolution detected from each step. **(B)** Genetic exchange test for FadL protein families. The genetic distance among *E. coli* strains within each lineage was shown and compared to that among strains from different lineages, respectively. Mann-Whitney *U* tests were performed, and *p*-values larger than 0.05 are highlighted in red. The significant *p*-values but with larger intra-lineage distance are shown in blue. A-A, B1-B1, B2-B2, D1-D1, D2-D2, or E-E represent strain pairs from the same lineage: A, B1, B2, D1, D2, or E, respectively. “x-y” represents pairs of strains from different lineages, e.g., A-B1, A-B2, D1-D2, etc. The *p*-values for core-proteome (CP_*p*) were shown in the bottom. **(C)** Clustering of FadL proteins with the neighbor-joining tree. The robustness of the phylogenetic tree was examined by bootstrapping tests with 1,000 replicates, and the scores are indicated for nodes of no lower than 50%. The strains from the same *E. coli*, *Shigella* or *E. fergusonii* lineage are labeled with a unique colored sign. The diagram for core-proteome (CP) tree referred to [30] and was also shown. **(D)** Distribution of three indices reflecting local variation property of the FadL protein family in *E. coli*. *HVRanalysis* was used for the analysis with the window width and step size set as 5 and 1, respectively. The significant highly variable sites (HVS) were shown in red circles.

We took *fadL* as an example to demonstrate the mosaic evolution (Fig. 1B-C). For some lineages, for example, lineages D1 and E, the protein sequences of *fadL* gene showed no significant difference between the within-lineage distance and the inter-lineage distance (Fig. 1B). In some cases such as the lineage B2, the FadL protein sequences even showed significantly larger distance among strains within the same lineage than the inter-lineage distance (Fig. 1B). As control, however, the distance of *E. coli* core proteome was significantly larger between lineages than within the same lineages expect for D1, which showed no significant difference (Fig. 1B). The FadL protein tree showed apparent incongruence with the *E. coli* lineage tree based on the core proteome, featured by the divergence and cluster separation of proteins from the strains of the same lineages (e.g., MG1655 and ATCC8739 of lineage A, S88 and ED1a of lineage B2, etc.), or the convergence of proteins from the strains of different lineages (e.g., ATCC8739 of lineage A and ED1a of lineage B2) (Fig. 1C). The genetic exchange analysis and phylogenetic analysis on fadL nucleotide sequences showed consistence with the results on the protein sequences. Taken together, the results demonstrated that *E. coli fadL* gene experienced events of ingenic recombinations.

We futher proposed three indices, namely MutIndex1, MutIndex2 and MutIndex, to illustrate intuitively the property of local mutations of the proteins under mosaic evolution, to test the unevenness of mutation along the protein sequence, and to identify HVRs and the sequenc types of each HVR (Materials and Methods). With this strategy, among the 27 E. coli outer membrane protein families, 21 demonstrated striking mosaic evolution patterns (Fig. 1A).

For each protein, the three indices showed similar distribution patterns along the protein sequnence, all of which were significantly different from random distributions, confirming the mosaic evolution (Fig. 1D, FadL as an example). According to the distribution patterns of the indices, especially the composite index MutIndex, local fragments (HVRs) could be delineated, which were strikingly larger than the average values (Fig. 1D).

### 2. Overall conservation of *E. coli* OMP families under mosaic evolution

Most (14/21) of the candidate *E. coli* OMPs under mosaic evolution encode porins (Fig. 2A), including the known families of FhuA, LamB, OmpA, OmpC and OmpF [29, 30]. Some flagella or pilus components (4/21), adhesins or antigens (2/21) and factors functioning on the membrane (1/21) also show the patterns of mosaic evolution (Fig. 2A). A majority of the genes have essential functions for bacterial cells and are conserved among *E. coli* lineages and strains, execept for *ag43*, which showed poor conservation (Fig. 2A). As surface antigen, the exact function and essentiality of Ag43 in E. coli cells remain unclear. For each of the other OMP families detected with mosaic evolution, each E. coli strain contains no more than one gene copy, and the genetic loci show good collinearity among different E. coli lineages (Fig. 2B). The *ag43* loci also show apparent collinearity among the strains bearing the gene. Therefore, the OMP genes under mosaic evolution are generally conserved.

**Figure 2.**
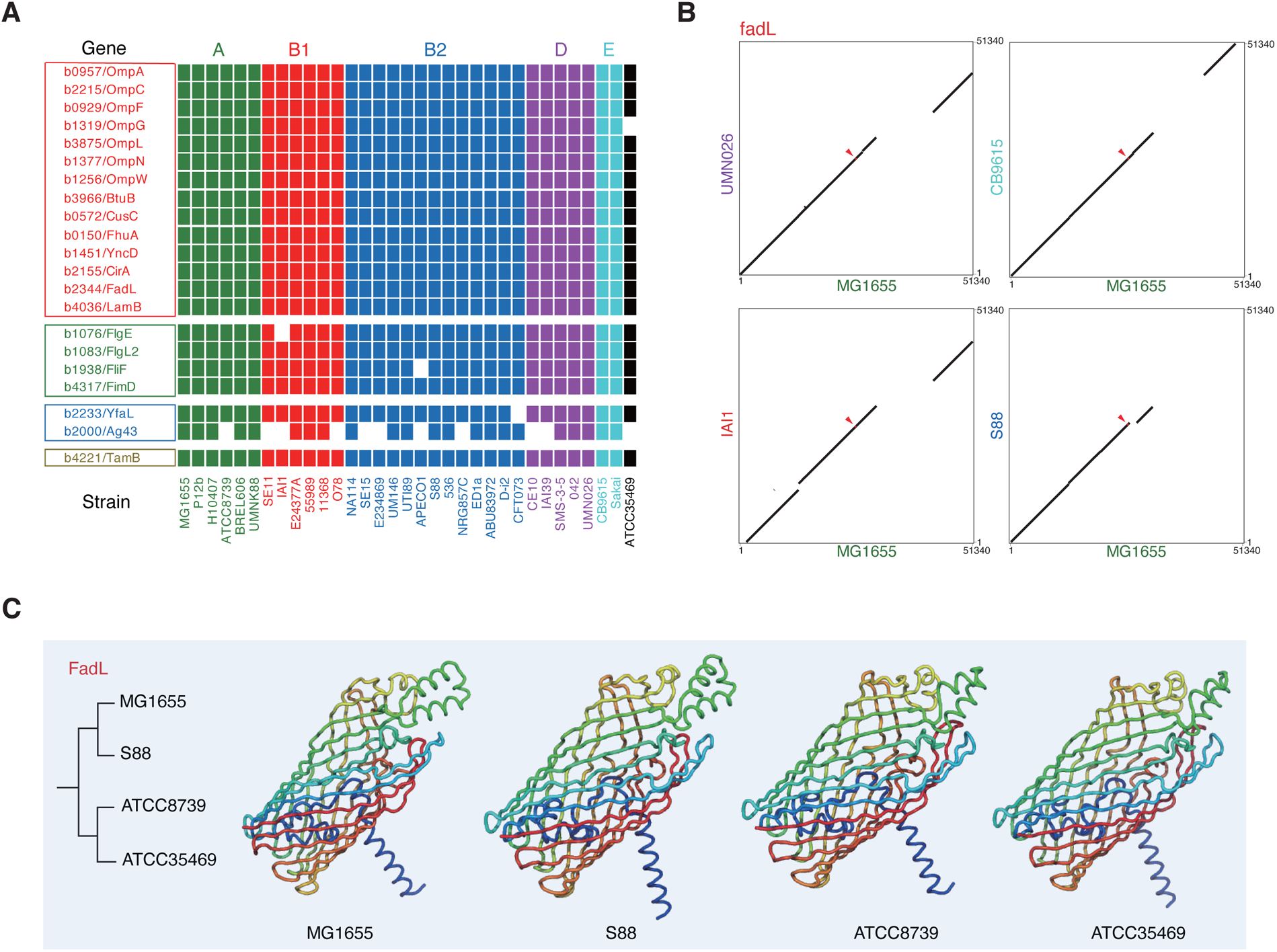
Conservation of E. coli OMPs under mosaic evolution. **(A)** Distribution of OMPs under mosaic evolution in *E. coli* strains. Missing of a gene in a strain was shown in white while the presence was shown in different colors for different lineages. Genes were grouped according to their main functions, including beta-barrel proteins, flagella and fimbriae components, adhesins and membrane-attaching antigens, and other transmembrane functional factors. **(B)** Collinearity of *fadL* gene and the flanking nucleotide sequences (25 kb at each side) between *E. coli* strains from different lineages. **(C)** The tertiary structures of FadL proteins of representative E. coli strains from different sequence clusters.

Tertiary structure of the proteins was also modeled and compared. Generally, the structure of full-length proteins appeared conserved. Within the same family, the proteins of different full-length sequence types or from different phylogenetic clusters always showed similar overall structure, implying that the basic function of the proteins were not likely disrupted by the local recombinations (Fig. 2C).

### 3. Transmembrane topology and structure of the *E. coli* OMP HVRs

The transmembrane topology was predicted with DeepTMHMM on the OMPs under mosaic evolution. As expected, the HVRs often coincided with the extracellular fragments for these proteins. For example, all the five HVRs of FadL are located in the extracellular regions predicted by DeepTMHMM, and the HVRs of OmpN are also outside of the cells (Fig. 3A). Similarly, for many other proteins, such as OmpG, OmpL, OmpW and others, also showed overlapping between all or partial HVRs and the extracellular protein fragments.

**Figure 3.**
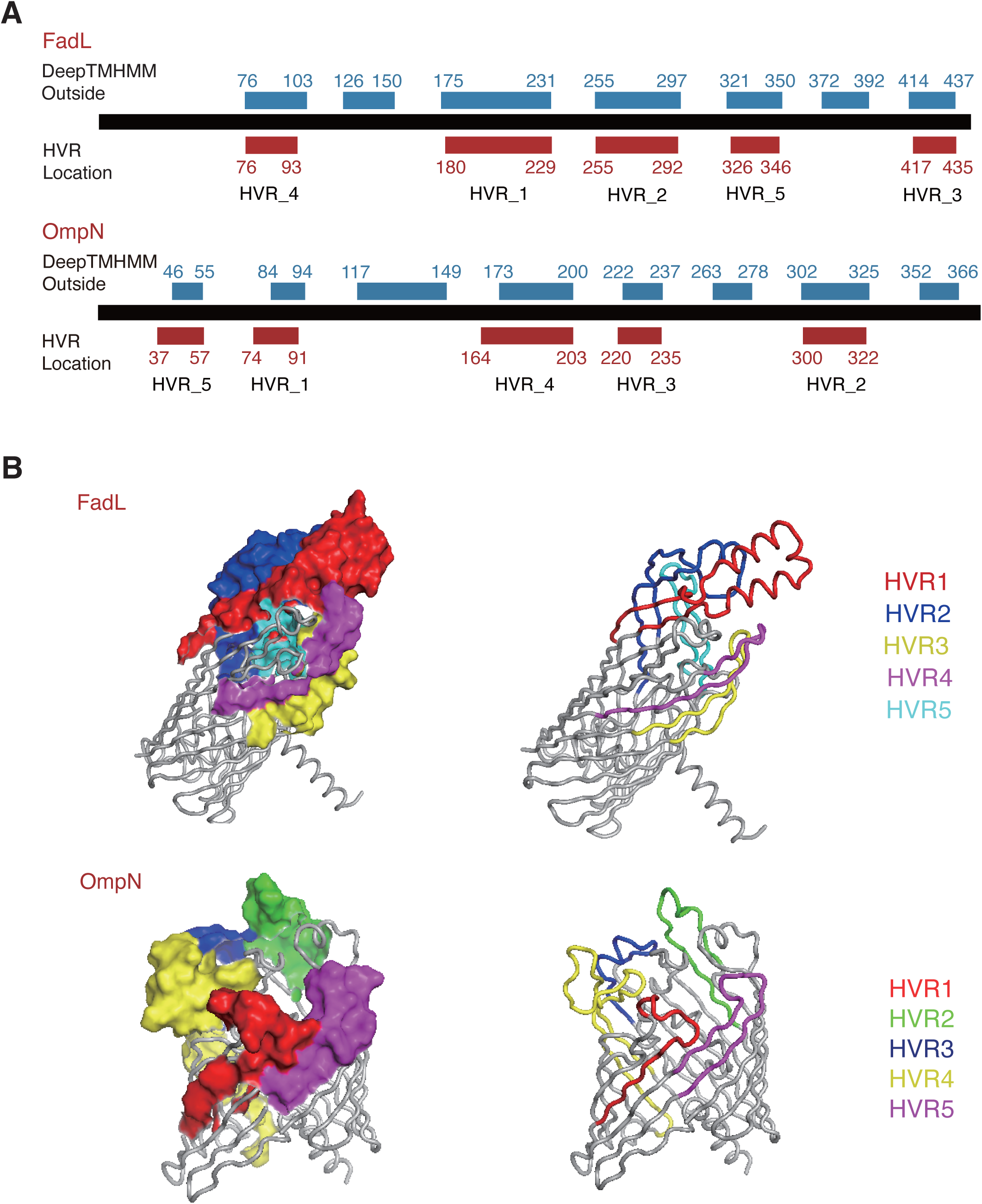
Transmembrane topology and tertiary structure of *E. coli* FadL and OmpN HVRs. **(A)** Transmembrane topology diagrams and the location of HVRs for FadL and OmpN. **(B)** The tertiary structure of FadL and OmpN, and the location of HVRs. Both surface and cartoon modes were shown for the HVRs. The *E. coli* MG1655 proteins were used for the analysis.

Consistent with the topology, the tertiary structure modeled for these proteins also suggested the frequent extracellular exposure of the HVRs. For example, all the HVRs of FadL are located in the surface of extracellular loops (Fig. 3B). Similarly, the HVRs of OmpN are also located in the protruding regions of the extracellular protein loops (Fig. 3B). Many of these proteins, including FadL, OmpW, and even flagella-related proteins, have been reported to serve as binding receptors of various bacteriophages. Although the exact interactions are unclear on these proteins, previous observations on FhuA, LamB, OmpA, OmpC and OmpF suggest the HVRs to be potentially important phage-binding sites.

### 4. Limited sequence types of the OMP HVRs in *E. coli* strains

According to the definition, OMP-encoding genes under mosaic evolution should experience local recombinations more often for the HVRs, and the sequence types for most HVRs are hypothesized to be with a limited number. We therefore selected some of the protein families, resolved the HVRs and observed the distribution of HVR sequence types in the correponding proteins among an enlarged size of *E. coli* strains. LamB and FadL were selected as examples that have one and multiple HVRs respectively.

LamB was detected from 3073 (99%) out of the total 3104 representative *E. coli* genomes (Fig. 4A). One HVR (HVR1) and four major sequence types (HVR1_1 ∼ 4) were detected from the 40 *E. coli* or close strains used for mosaic evolution screening in this study, while each major HVR sequence type contains 1 to 2 subtypes with 1-3 amino acid substitutions (Fig. 4B). Among the 3073 LamB-positive strains, no any new major HVR sequence type was detected (Fig. 4B). 96.9% (2978/3073) of the strains showed the HVR sequence sybtypes exactly consistent with the known ones, while only 2.8% (87/3073) and 0.3% (8/3073) showed 1-2 and 3 amino acid substitutions compared to the known sequence subtypes, respectively (Fig. 4B).

**Figure 4.**
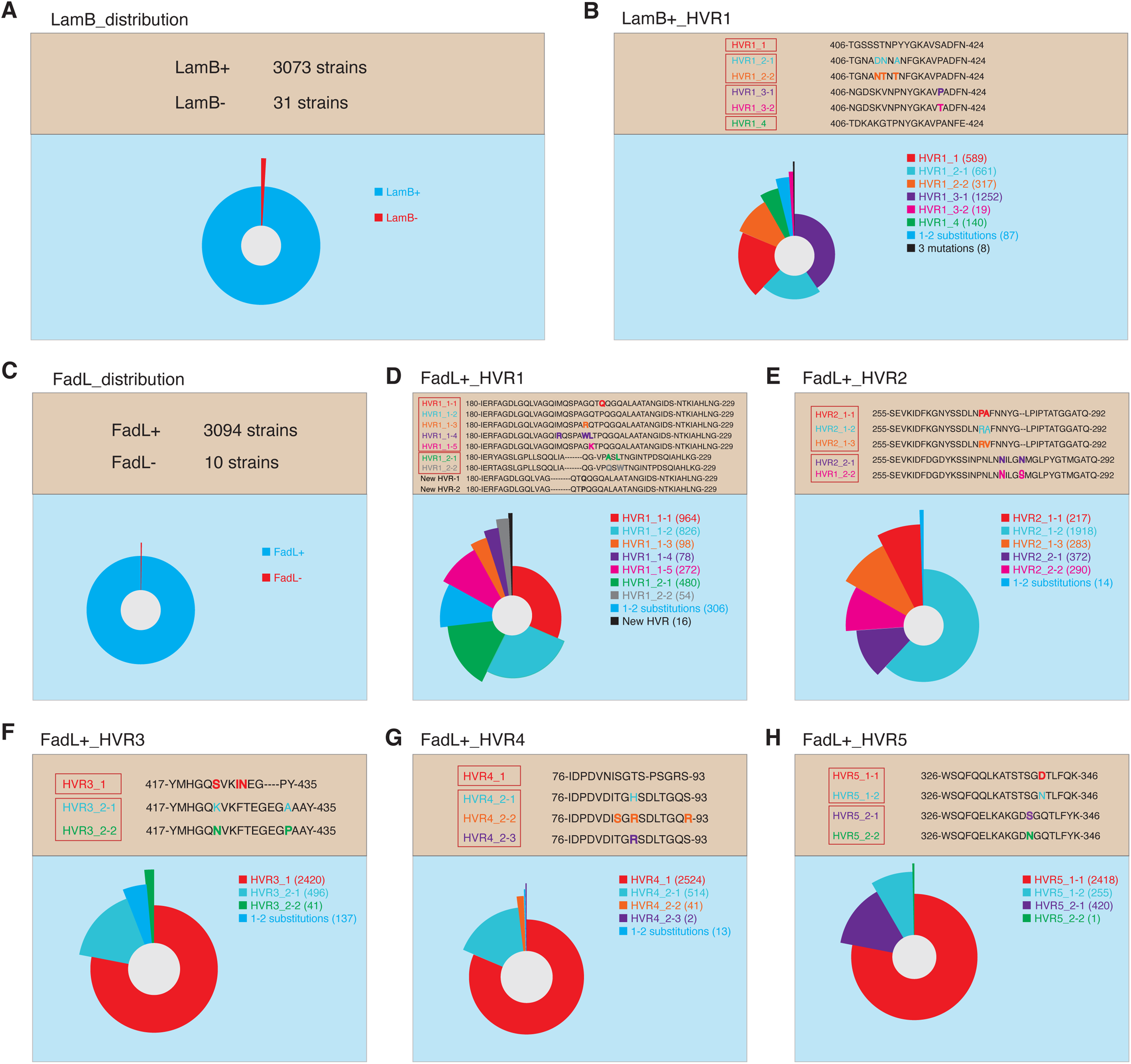
Distribution of LamB and FadL HVR sequence types in *E. coli* strains. **(A)** Distribution of LamB in 3104 *E. coli* strains. **(B)** Distribution of known LamB HVR sequence types and subtypes in the independent genome set of 3073 LamB-containing *E. coli* strains. **(C)** Distribution of FadL in 3104 *E. coli* strains. **(D)** - **(H)** Distribution of known FadL HVR sequence types and subtypes in the independent genome set of 3094 FadL-containing *E. coli* strains.

Nearly 99.7% (3094/3104) of the *E. coli* genomes could be detected with the FadL-encoding gene (Fig. 4C). For the 5 HVRs of FadL proteins, HVR1∼5 each contains two major sequence types, and 7, 5, 3, 4 and 4 subtypes respectively (Fig. 4D-H). Except for HVR1, no novel major sequence type was detected from the 3094 FadL-positive *E. coli* strains for each HVR (Fig. 4D-H). A new major sequence type was detected for HVR1 from 16 strains, including two sequence subtypes with only one amino acid substitution (Fig. 4D). For HVR1∼5, there are 89.6% (2272/3094), 99.5% (3080/3094), 95.6% (2957/3094), 99.6% (3081/3094) and 100% (3094/3094) of the strains showing sequence subtypes exactly same with the known ones (Fig. 4D-H). No subtype was detected from each HVR showing more than 2 amino acid substitutions compared to known sequence subtypes (Fig. 4D-H).

Taken together, the results demonstrated that the number of major sequence types for the HVRs of OMPs under mosaic evolution is very limited. Natural mutations also happen, but slowly within the HVRs.

### 5. Micro-evolution of the HVRs of *E. coli* OMPs under mosaic evolution

We calculated the selection pressure for each major sequence type of the HVRs of FadL in *E. coli*. Despite the non-positive selection for the whole fragments of the HVRs and most codon positions, several sites were detected with significantly positive selection (Fig. 5A-E). Specifically, the 19^th^, 22^nd^, 23^rd^ and 25^th^ codons of HVR1_1, the 4^th^, 19^th^, 23^rd^, 24^th^ and 25^th^ codons of HVR1_2, and the 13^th^ codon of HVR5_2 were all positively selected (Fig. 5A, 5E, 5F). Previously, we also detected a few sites under significant positive selection within LamB and and OmpC HVRs [30]. Therefore, the HVRs for individual sequence types could also experience positive selection, albeit slowly, so as to make sequence varing, generating new major HVR sequence types eventually.

**Figure 5.**
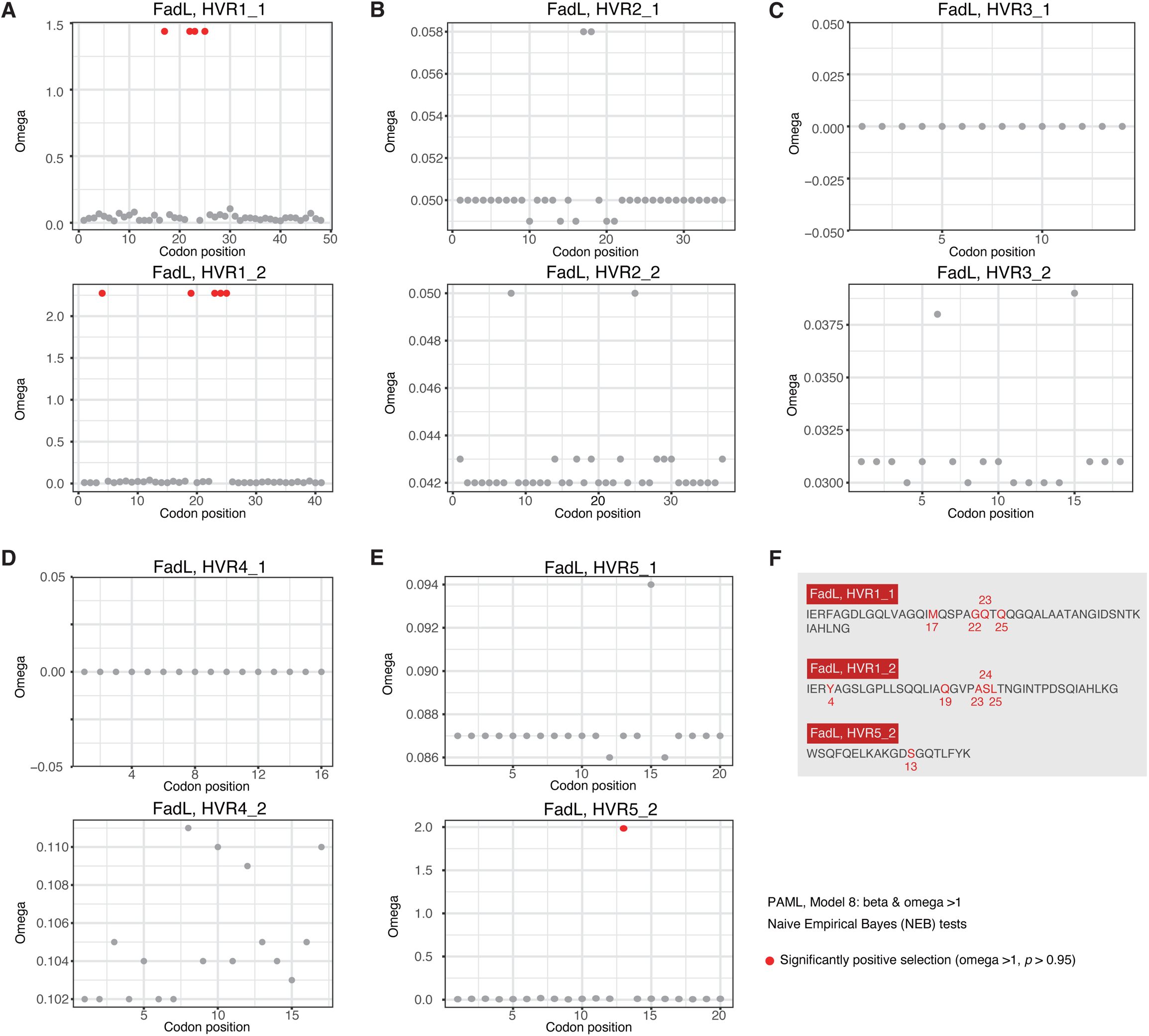
Selection pressure of FadL HVRs in *E. coli*. **(A)** - **(E)** Omega distribution along the sites of each major HVR sequence type of FadL in *E. coli*. The sites of significant positive selection were shown in red. **(F)** Summary of the positively selected sites detected in *E. coli* FadL HVRs.

### 6. Immunogenicity of the HVRs of *E. coli* OMPs under mosaic evolution

The HVRs could be good antigens due to their extracellular localization, hydrophilicity and relative conservation of the sequence types. Consequently, we tested their potential immunogenicity. Both LamB and FadL were selected as representative proteins for analysis.

LamB of the each HVR sequence type was predicted to have high antigenicity with a ANTIGENpro probability of >0.88 (Fig. 6A). The antigenicity of control LamB proteins with HVRs removed (LamB_noHVR) was reduced apparently compared to the wild-type LamB proteins (LamB) or LamB proteins removed of the same length of non-HVR fragments (LamB_len_ctrl), suggesting the important contribution of HVRs to the antigenicity of LamB proteins (Fig. 6A). Linear B-cell epitopes were further predicted from LamB proteins with an online tool ABCpred and three newly developed tools with different features and algorithms, B_MM, BPBB and JBFB (Materials and Methods). With ABCpred, we detected a significant B-cell epitope, which is well consistent with the HVR of LamB (Fig. 6B). It was noted that, in all the four LamB proteins with different HVR sequence types, the most significant B-cell epitopes are with the same location in the HVRs (Fig. 6B). There is also another epitope with large score and high rank within the HVR in each representative LamB protein (Fig. 6B). Consistent with ABCpred, B_MM, BPBB and JBFB all predicted that the HVR of MG1655 showed the strongest immunogenicity along the LamB protein (Fig. 6C, red). The LamB proteins of different HVR sequence types showed the simiar results (Fig. 6D). Furthermore, DiscoTope-2.0 and CBTOPE were used for prediction of structural B-cell epitopes. Consistent with the observation for linear epitopes, both tools predicted significant structural B-cell epitopes from LamB proteins of different sequence types, all being located in the HVRs (Fig. 6E). Simiar observations were obtained from FadL. FadL from different sequence clusters also show good antigenicity (Fig. 6F), and the HVRs exhibit strong immunogenicity (Fig. 6G).

**Figure 6.**
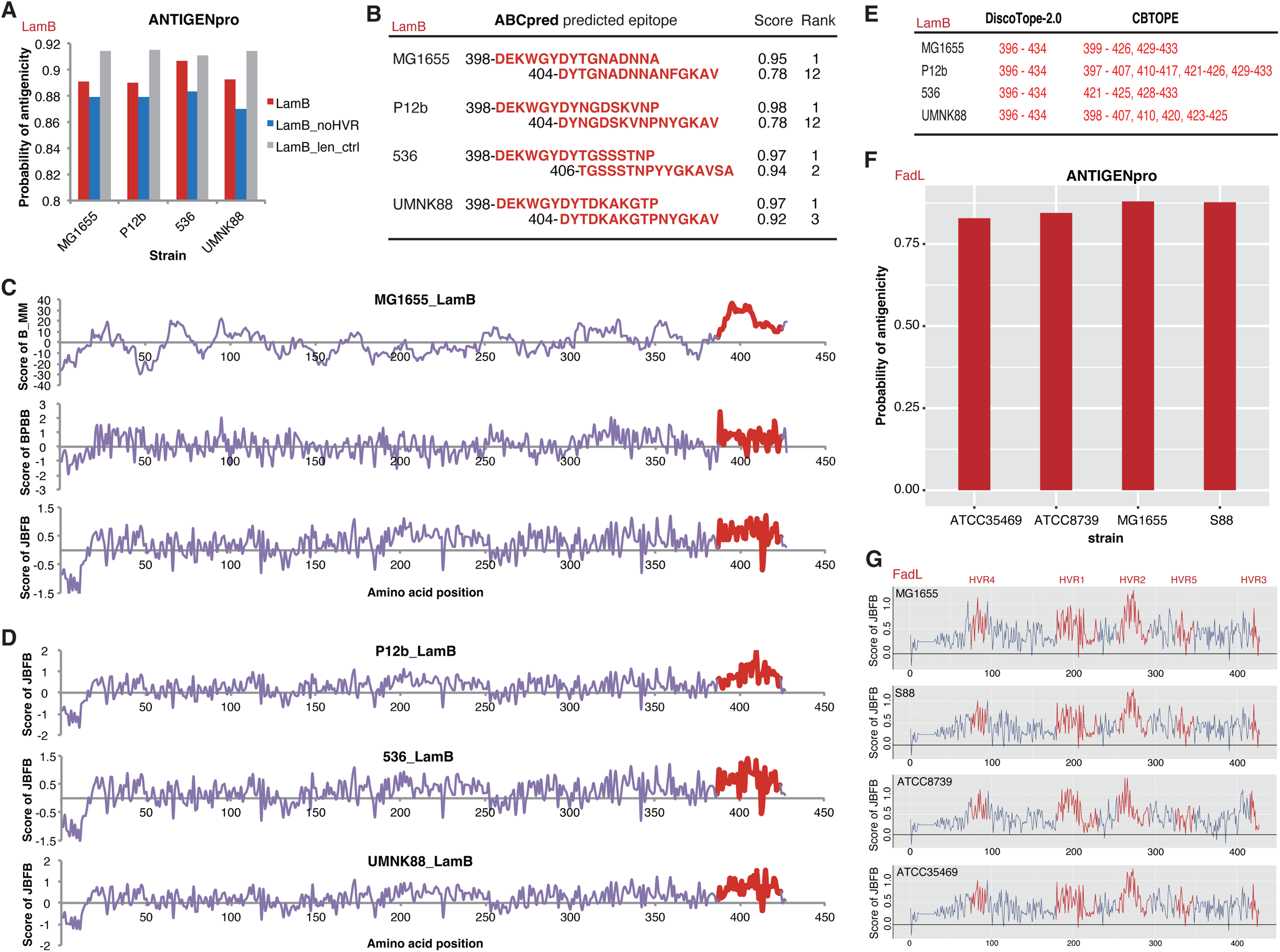
Immunogenicity of *E. coli* LamB and FadL HVRs. **(A)** Antigenicity of LamB of different sequence clusters. LamB, LamB_noHVR and LamB_len_ctrl represented the wild-type LamB, HVR-removed LamB, and LamB removed of non-HVR fragments with the total length identical to that of the HVRs, respectively. **(B)** The scores and ranks of ABCpred predicted B-cell epitopes in LamB HVRs. **(C)** The predicted scores of B_MM, BPBB and JBFB on the linear B-cell epitopes in MG1655 LamB. The scores for the HVRs were highlighted in red. **(D)** The predicted scores of JBFB on the linear B-cell epitopes in LamB proteins from different sequence clusters. **(E)** Structural B-cell epitopes in LamB HVRs predicted by DiscoTope-2.0 and CBTOPE. **(F)** Antigenicity of FadL of different sequence clusters. **(D)** The predicted scores of JBFB on the linear B-cell epitopes in FadL proteins from different sequence clusters. The scores for the HVRs were highlighted in red.

Taken together, the results suggested that the OMPs under mosaic evolution could potentially be used for antibody development. The HVRs of various sequence types could serve as targetted antigens, which often bear strongly immunogenic epitopes.

## Discussion

Several OMP families have been reported to show patterns of mosaic evolution in different bacteria [29, 30, 32–35]. However, it is unknown how many OMP families experiencing such type of evolution in a bacterial species. To our knowledge, this study made the first genome-wide investigation on the mosaic evolution in OMP-encoding genes in *E. coli*. To our surprise, around a half of the OMP genes in *E. coli* can be detected with ingenic recombination events, as predicted by RDP4. Many of the recombinations generate no or only atypical changes in protein level. However, there remain 21 OMP families showing typical patterns of mosaic evolution, including the incongruence of gene or protein trees featured by inter-lineage convergence and clustering, larger within-lineage but smaller inter-lineage molecular distance, and being detected with significant uneven fragmental mutation rates and typical HVRs. The five known porins under mosaic evolution are all included in the 21 candidate proteins. Therefore, the real number of bacterial OMPs under mosaic evolution could be much larger than we expected. It would be interesting to investigate whether it is the case in other bacteria and even in Gram-positive bacteria. We also noticed that a large majority of the OMPs under mosaic evolution are beta-barrel proteins. It is likely that mosaic evolution widely happens in genes encoding bacterial beta-barrel proteins.

Most of the OMPs under mosaic evolution have functions essential for bacterial survival. The proteins from different sequence clusters always show conserved tertiary structures in general (Fig. 2C). With very few exceptions, the genes are present in and genomically conserved among *E. coli* strains (Fig. 2A-B). The proteins are often receptors of phages, and the HVRs are important phage-binding sites. In this research, we also found that the HVRs are also candidate fragments that could elicit strong B-cell immune responses (Fig. 6). Therefore, the theory on bacterial mosaic evolution was further consolidated by this study. Bacteria could rapidly escape from the attacks of harmful factors, such as phages, immune factors and others, and maintain the essential function by rapid ingenic recombination for the HVRs [30]. These HVRs have actually been experiencing a long period of evolution and selection and diverged from their ancestral sequence type very slowly, and therefore the fragmental conversion do not lead to bacterial fitness problems. In fact, within the major HVR sequence type, we did find, though very few, HVR sites under positive selection, in FadL (Fig. 5), LamB and OmpC [30]. There remains a question for the theory as to the sources of HVRs. It was reported that various recombinations and gene conversions are widely spread in *E. coli* and other bacteria [50, 51], although the recomnination sources are also unknown. The microbiota co-present in the environment where the target strain was isolated could serve as an optional library for the genetic molecule sources.

The study also paved the foundation for potential applications of the mosaic evolution and HVRs for development of anti-bacterial reagents, for example, monoclonal therapeutic antibodies. Traditionally, the conserved peptide fragments rather than the HVRs, should be considered as stable antigen candidates for antibody development. However, the long-term co-evolution between bacteria and the hosts and the selection have often made the conserved peptide fragments not located extracellularly, unexposed and lowly immunogenic. The HVRs are located extracellularly, accessible by the immune factors, and with strong imminogenicity. The essential function of the proteins and the conserved presence among strains reduce the risk of off-target. The recombination nature of HVRs ensures the limited number of major sequence types and the predictability (Fig. 4), making it possible to develop a mixed pool of antibodies against each major HVR sequence types. Each HVR sequence type or subtype seemed to show similarly strong immunogenicity (Fig. 6). These results all suggest the promising application of these HVRs [52]. However, the practical effect should be observed with more elaborate and extensive experimental validation. Specifically, it would be important to test whether an antibody against a HVR subtype could cross-react with other subtypes of the same major sequence type with 1-2 amino acid variations.

## Materials and Methods

### 1. Bacterial species and strains

The previously annotated 39 representative *E. coli* or *Shigella* strains and one outgroup strain, *E. fergusonii* ATCC35469, were used in this study for screening of OMPs under mosaic evolution [29, 30]. To observe the distribution of HVR sequence types, 3104 *E. coli* strains whose genomes were fully sequenced, assembled and well annotated in the NCBI Genome database were selected, and their genome sequences and annotation files were downloaded (Supplementary Dataset 1).

### 2. Identification of *E. coli* OMP families

OMPs were predicted from the genome-derived proteins of *E. coli* MG1655 using PRED-TMBB [36] and TMBETADISC-RBF [37] with default settings. The intersection of positive predictions from the two tools were considered as the OMPs, with negative predictions but putative OMPs included, and positive predictions but putative non-OMPs excluded. Furthermore, BLASTp was implmented to search the homologs of the MG1655 OMPs in the other 39 *Escherichia* and *Shigella* strains. The paratemers were set as ≥ 80% identity for an average length coverage of ≥ 90%.

### 3. Screening of OMP families under mosaic evolution

For each OMP family, the nucleotide sequences encoding the proteins were analyzed for possible ingenic recombination with RDP4 [38]. Ingenic recombination was defined if only at least one significant recombination event (*p* < 0.05) was identified by any one model or statistical approach integrated in RDP4. For each OMP family predicted by RDP4 with ingenic recombination, MEGA7.0 [39] was used to build the gene and protein trees with the neighbor-joining algorithm. The trees were manually checked and compared with the lineage tree based on the core proteome of the 40 *Escherichia* and *Shigella* strains, and thereby the OMP families were identified whose trees were apparently incongruent with the lineage tree. The core proteome and evolutionary analysis referred to previous reports [29, 30]. Furthermore, for the OMP families whose gene and protein trees both deviated from the lineage tree, genetic exchange tests were conducted at both nucleic acid and protein levels. The positive observation of genetic exchange tests was defined to an OMP family for which at least one lineage showed a change from significance to non-significance in the Wilcoxon rank-sum tests for within-lineage vs. inter-lineage genetic distance, or a reversal in direction at both nucleic acid and protein levels [30].

### 4. Validation of mosaic evolution and HVR identificaiton

A Python package, *HVRanalysis*, was developed to validate mosaic evolution and to identify the composition and sequence types of HVRs directly. We proposed three indices, MutIndex1, MutIndex2 and MutIndex, to comprehensively measure the local variation characteristics of proteins. In the first place, the protein sequences of an interesting family are aligned using CLUSTALW [40] with the default settings. Given *n* proteins, each with an aligned length *L*, peptide segments are continuously extracted by sliding a predefined window of width *l* and step size *s* across the aligned sequence of each protein simultaneously. For each set of *n* peptide segments obtained after each step, the number of sequence types *m* and the specific sequence types denoted as *S_1_*, *S*_2_, …, *S_m_*, are analyzed after redundancy removal. The frequency *F_Si_* of each sequence type *S_i_* (*i* = *1*, *2*, …, *m*) among the *n* peptide segments is calculated. In each window, the *n* peptide segments are observed site by site. A variation site is recorded whenever a different amino acid (or deletion) is found at the corresponding site in any sequence. The variant sites *v* are counted among the window length *l*.

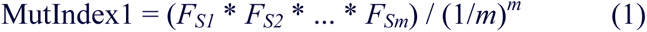

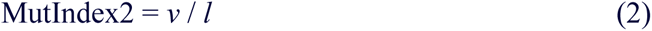

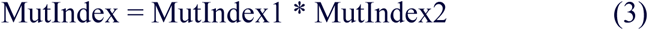

For each set of segments, MutIndex1, MutIndex2 and MutIndex are calculated according to the above formulas. MutIndex1 reflects the distribution balance of different sequence types within local regions within the strain population, MutIndex2 reflects the frequency of variations within local regions, and MutIndex combines MutIndex1 and MutIndex2 to comprehensively reflect the distribution of sequence types and variation sites. For each index, the mean and variance are calculated across the windows along the full protein alignment length, and a Gaussian distribution is simulated. A Kolmogorov-Smirnov (KS) test is performed to assess whether the distribution of the actual index conforms to or deviates from the simulated Gaussian distribution. If the KS test *p*-value is below the predefined significance level of 0.05, the corresponding index is not considered to follow a normal distribution along the protein alignment length, indicating the presence of local mosaicism. Continuous sites outside the 95% confidence interval of the simulated normal distribution are extracted, and the segments they compose are recorded as HVRs. The amino acid sequences of each HVR are extracted from the strains and deduplicated to obtain HVR sequence type or subtypes.

*HVRanalysis* is freely accessible at http://61.160.194.165:3080/HVRanalysis. In this study, the default settings are used for *HVRanalysis* with *l* = 10 and *s* = 1 if not specified.

### 5. Comparative genomic analysis

Each OMP under mosaic evolution is aligned against the proteins encoded by the genomes of various strains using BLASTp with identity >= 70% over an average length coverage of >= 70%, to analyze the copy number of homologous genes in corresponding strains. The gene loci are delinated within the genome, and continuous nucleotide sequences including the gene sequences and the 25K flanking sequences at both upstream and downstream are retrieved. PipMaker [41] is used to perform collinearity analysis among the genes and their flanking sequences from different strains.

### 6. Distribution analysis of HVR sequence types among *E. coli* population

The OMPs under mosaic evolution identified in 40 *Escherichia* and *Shigella* strains (termed as reference strains and OMPs) are used to determine and label their respective HVRs and sequence types based on the results of *HVRanalysis*. The full-length protein sequences are aligned with BLASTp against the full proteome of 3104 *E. coli* strains downloaded from NCBI Genome. For alignments with 100% identity across the full length, the corresponding *E. coli* strains, their best-matched reference OMPs, and the HVR sequence types and subtypes are annotated. For the orthologous OMPs of the remaining *E. coli* strains, an in-house Python script is prepared to search each known HVR sequence subtype so as to annotate the HVRs, HVR sequence types and subtypes for the OMP orthologs in more strains. BLASTp is applied again to make mutual alignment for the unannotated OMP orthologs, followed by clustering the 100% identical ones. One representative protein is randomly selected from a cluster, multiple sequence alignments are performed to these representative OMPs and the reference OMPs, and the HVRs are annotated with new sequence types and subtypes identified according to the extent and patterns of variations.

### 7. Micro-evolution analysis of the HVRs

Based on sequence homology, each HVR was classified into major sequence types (with > 3 site mutations between each other). For a specific major HVR sequence type, the nucleic acid sequences were extracted from the corresponding OMPs of the reference *E. coli* strains, and selection analysis was performed using PAML [42]. The Model 8 was used, omega was estimated for each codon site, and Naive Empirical Bayes (NEB) tests were performed to assess positive selection, that is, omega > 1. The significance was preset as *p* > 0.95.

### 8. Transmembrane topology and structural analysis of OMPs and HVRs

DeepTMHMM [43] was used to predict the transmembrane topology of OMPs, while Phyre2 [44] and RoseTTAFold [45] were used to predict their tertiary structures. The topological and structural characteristics of the regions where HVRs are located were annotated.

### 9. Immunogenicity analysis of HVRs

ANTIGENpro [46] was used to analyze the antigenicity of the proteins representing each HVR sequence type. DiscoTope-2.0 [47] and CBTOPE [48] were used to analyze the structural B-cell epitopes. ABCpred [49] was used to analyze the linear B-cell epitopes. Besides, we also designed three new models, B_MM, BPBB and JBFB, to predict linear B-cell epitopes based on different features and algorithms. The models were also used for antigen epitope analysis on the OMPs of different HVR sequence types. The webserver, standalone models and the documents are accessible freely through the link: http://61.160.194.165:3080/B_pred.

## Authors’ Contribution

Y.W., A.P.W., W.K. and G.Z. conceived the project. X.C., C.C., Z.C., Y.Z., and Y.H. collected the data. X.C., C.C., Z.C., J.L., Y.Z., J.W., Z.L., G.H., and Y.W. performed the analysis. Y.Z., X.C., Z.C., J.L., Z.L., Y.H., and Y.W. designed and developed the bioinformatic methods. X.C., C.C., Z.C., J.L., Z.L., G.H., and Y.W. wrote the first draft. G.Z., A.P.W., W.K., and Y.W. revised the manuscript. All the authors approved the final version of manuscript.

## Supplemental Materials

**Supplemental Dataset 1. Information of *E. coli* strains with full genome sequences downloaded from the NCBI Genome database.**

